# Salmonid chromosome evolution as revealed by a novel method for comparing RADseq linkage maps

**DOI:** 10.1101/039164

**Authors:** Ben J. G. Sutherland, Thierry Gosselin, Eric Normandeau, Manuel Lamothe, Nathalie Isabel, Céline Audet, Louis Bernatchez

## Abstract

Whole genome duplication (WGD) can provide material for evolutionary innovation. Assembly of large, outbred eukaryotic genomes can be difficult, but structural rearrangements within such taxa can be investigated using linkage maps. RAD sequencing provides unprecedented ability to generate high-density linkage maps for non-model species, but can result in low numbers of homologous markers between species due to phylogenetic distance or differences in library preparation. Family Salmonidae is ideal for studying the effects of WGD as the ancestral salmonid underwent WGD relatively recently, around 65 million years ago, then rediploidized and diversified. Extensive synteny between orthologous chromosomes occurs in extant salmonids, but each species has both conserved and unique chromosome arm fusions and fissions. Here we generate a high-density linkage map (3826 markers) for the *Salvelinus* genera (Brook Charr *S. fontinalis*), and then identify orthologous chromosome arms among the other available salmonid high-density linkage maps, including six species of *Oncorhynchus*, and one species for each of *Salmo* and *Coregonus*, as well as the sister group for the salmonids, *Esox lucius* for homeolog designation. To this end, we developed MapComp, a program that identifies identical and proximal markers between linkage maps using a reference genome of a related species as an intermediate. This approach increases the number of comparable markers between linkage maps by 5-fold, enabling a characterization of the most likely history of retained chromosomal rearrangements post-WGD, and identifying several conserved chromosomal inversions. Analyses of RADseq-based linkage maps from other taxa will also benefit from MapComp, available at: https://github.com/enormandeau/mapcomp/

## Introduction

Whole genome duplication (WGD) can provide the raw material for evolutionary innovation by generating redundant copies of all chromosomes (i.e. producing homeologous chromosome pairs). After WGD, the genome can then undergo rediploidization while retaining all homeologous chromosomes, thereby doubling the pre-duplication chromosome number. After rediploidization, homeologous pairs diverge through several mechanisms. Gene copies can evolve new functions, sub-functionalize the original function between the two copies or, most frequently, accumulate mutations that disrupt functionality of one copy (Ohno 1970; Force et al. 1999; Brunet et al. 2006). Cross-taxa analyses suggest that during rediploidization, genes that only keep one copy are often retained within the same homeologous chromosome (Sankoff et al. 2010). However, this non-random retention is not always observed (Berthelot et al. 2014). Interestingly, rediploidization does not always complete. For example, some homeologous chromosome arms in the salmonids continue recombining, which is referred to as residual tetrasomy (Allendorf et al. 2015).

Eukaryotic genomes with ancestral WGD (e.g. pseudotetraploid genomes) are challenging to assemble (Davidson et al. 2010). Linkage maps can be highly useful for comparing chromosomal evolution among lineages using shared markers between maps to identify corresponding chromosomes (i.e. orthologous chromosomes) between species (Naish et al. 2013; Kodama et al. 2014). Furthermore, high quality, dense linkage maps are valuable for validating and orienting genomic scaffolds (Fierst 2015; Mascher & Stein 2014), especially for cases of residual polyploidy, large genome size, and high repeat content (Ming & Man Wai 2015; Amores et al. 2014). Recent advances in sequencing, such as through reduced-representation library sequencing (e.g. RADseq) (Baird et al. 2008; Elshire et al. 2011; Andrews et al. 2016), have made high-density linkage maps increasingly easy to produce. These methods provide thousands of markers without requiring marker design effort. They can also generate haplotype loci (i.e. loci with more than one SNP) to improve mapping resolution by increasing ability to assign alleles to a parent (Catchen et al. 2011). RADseq-based SNP markers are contained in short sequence fragments, which allow for mapping against a genome to identify nearby genes or physical distances between markers (Amores et al. 2011; Henning et al. 2014). RADseq also enables comparative genomics through the use of direct marker-to-marker comparisons to find homologous (i.e. nearly identical markers between species) markers between linkage maps (Kodama et al. 2014). The ability to identify orthologous chromosomes between maps is dependent on being able to identify enough shared markers between species, and this decreases with phylogenetic distance due to sequence divergence (Gonen et al. 2015). This issue is compounded further when different protocols or restriction enzymes are used for library generation. Due to this, it has been suggested to use a common enzyme and protocol to ensure compatibility of maps (Larson et al. 2015), but this may not always be possible or desirable. Here we developed a method to use an intermediate reference genome for integrating linkage maps of different species by pairing both homologous and proximal markers in order to investigate orthologous and syntenic relationships among species.

Salmonids are a highly relevant study system for investigating the effects of WGD. The ancestor of modern day salmonids experienced a relatively recent salmonid-specific (4R) WGD, and subsequently underwent rediploidization (Allendorf & Thorgaard 1984; Davidson et al. 2010). Post-WGD, the salmonid lineage diversified into three subfamilies, 11 genera and more than 60 described species (Crête-Lafrenière et al. 2012), although this diversification was likely due to environmental factors rather than being caused by WGD (Macqueen & Johnston 2014). Analysis of the Atlantic Salmon genome suggests that the rediploidization process was rapid and that two classes of homeolog similarity exist: immediately rediploidized homeologs and those remaining in residual tetrasomy (see Figure 3b in Lien et al. 2016). Although much remains to be understood about this process in salmonids, fundamental work on chromosomal evolution has been conducted using cytogenetics and genetic maps (Phillips & Ráb 2001; Naish et al. 2013). From linkage map comparisons using homologous markers, it is known that the same eight pairs of orthologous homeologs are residually tetrasomic in Chinook Salmon (Brieuc et al. 2014), Coho Salmon (Kodama et al. 2014) and Sockeye Salmon (Larson et al. 2015).

Chromosomal evolution within family Salmonidae (i.e. whitefish, trout, charr and salmon) is typified by centric Robertsonian fusions, whereby two acrocentric chromosomes fuse into one larger metacentric chromosome, retaining the total number of chromosome arms *(nombre fondamental* (NF)=100) (Phillips & Ráb 2001). Fissions and whole arm translocations can also occur, subsequently separating the fused metacentric chromosomes. Cytogenetic research has identified the presence of two major karyotype groups in salmonids that differ in the number of retained chromosome fusion events. Type A species (2n=~80 chromosomes) have more acrocentric than metacentric chromosomes, whereas Type B species (2n=~60 chromosomes) have more metacentric than acrocentric chromosomes (Phillips & Ráb 2001). Adaptive mechanisms or selective forces driving these rearrangements and correlation with habitat or species biology remain generally unknown (Phillips & Ráb 2001). Among species, highly retained micro and macro synteny is expected between orthologous chromosome arms (Kodama et al. 2014), and within a species collinearity is observed between homeologs (Berthelot et al. 2014). Conservation of chromosome fusions has been partially explored between Chinook *O. tshawytscha* and Coho Salmon *O. kisutch* with Atlantic Salmon, allowing for the phylogenetic timing of rearrangements in these species (Kodama et al. 2014). However, this characterization has not been examined across the salmonid lineage using all available high-density maps. Furthermore, some genera do not yet have high-density genetic maps available.

The fusions of acrocentric to metacentric chromosomes may have important implications for the rediploidization process in salmonid evolution. Residual tetrasomy is thought to always involve at least one homeolog in a metacentric fusion (Wright et al. 1983; Allendorf et al. 2015), indicating that fusions can affect rediploidization efficiency. Metacentric chromosomes are more likely to form tetravalents at meiosis, permitting recombination between homeologs and preventing sequence divergence particularly in male salmonid telomeric regions (Allendorf & Danzmann 1997; Waples et al. 2016; Allendorf et al. 2015; May & Delany 2015). Salmonid species with few fusions may therefore provide new insight on the rediploidization process. This highlights the importance of characterizing orthologous relationships between chromosome arms and the acrocentric or metacentric states of chromosomes in order to gain information regarding expectations for levels of homeolog differentiation within each species.

High-density linkage maps have been constructed for Lake Whitefish *Coregonus clupeaformis* (Gagnaire et al. 2013), Atlantic Salmon *S. salar* (Lien et al. 2011; Gonen et al. 2014) and members of *Oncorhynchus* including Rainbow Trout *O. mykiss* (Miller et al. 2012; Palti et al. 2015), Chinook Salmon *O. tshawytscha* (Brieuc et al. 2014), Coho salmon *O. kisutch* (Kodama et al. 2014), Pink Salmon *O. gorbuscha* (Limborg et al. 2014), Chum Salmon *O. keta* (Waples et al. 2016) and Sockeye Salmon *O. nerka* (Everett et al. 2012; Larson et al. 2015). No high-density maps exist for members of *Salvelinus*, but low-density microsatellite-based maps exist for Arctic Charr *S. alpinus* and Brook Charr *S. fontinalis* (Woram et al. 2004; Timusk et al. 2011), as well as a low-density (~300 marker) EST-derived SNP map for *S. fontinalis* (Sauvage et al. 2012a). Genome assemblies exist for Rainbow Trout (Berthelot et al. 2014) and Atlantic Salmon (Lien et al. 2016). A genome assembly and low-density genetic map are also available for Northern Pike *Esox lucius*, a sister species to the salmonid WGD (Rondeau et al. 2014). With these resources available, it becomes especially valuable to integrate the information from all of the maps to detail the chromosomal evolution of the salmonids.

In this study, we use a mapping family previously used to generate a low-density EST-derived SNP linkage map (Sauvage et al. 2012a) to produce the first high-density RADseq map for the genus *Salvelinus*, the Brook Charr *S. fontinalis*. Brook Charr is a species of importance for conservation, aquaculture and fisheries, and an underrepresented lineage of Salmonidae in terms of genomic resource availability.

Further, we developed MapComp, a program to enable comparisons of genetic maps built from related species with or without the same RADseq protocol using an intermediate reference genome. MapComp follows earlier proposed approaches to integrate non-model maps with model species genomes (Sarropoulou et al. 2008). It identifies on average 5-fold more marker pairs between linkage maps than methods relying on identical markers only, and creates pairwise comparison plots for data visualization. MapComp enabled a detailed characterization of the orthologous and homeologous chromosome arms representing all main genera comprised within the salmonid family. This characterization enabled the identification of the most likely historical chromosomal rearrangements occurring at different levels of the salmonid phylogeny, including some potential inversion events. This comprehensive view provides new insight on the post-WGD chromosome evolution of Family Salmonidae.

## Materials and Methods

### Brook Charr genetic map

#### Animals

Full details regarding the experimental mapping family were reported previously (Sauvage et al. 2012a; 2012b). The F_0_ individuals were from a domestic population used in Québec aquaculture for 100 years, supplied here from the Pisciculture de la Jacques-Cartier (Cap-Sante, Quebec), and a wild anadromous population from Laval River (near Forestville, Québec) that have been kept in captivity for three generations at the Station aquicole de l’ISMER (Rimouski, Québec). Three biparental crosses of F_1_ individuals produced three F_2_ families, and the family with the largest number of surviving offspring was chosen to be the mapping family (n=192 full-sib F_2_ offspring).

#### DNA extraction, sampling preparation and sequencing

DNA was extracted from the fin of F_2_ offspring and F_1_ parents by high salt extraction (Aljanabi & Martinez 1997) with an additional RNase A digestion step (QIAGEN), as previously reported (Sauvage et al. 2012a). Quality of the extracted genomic DNA was quality validated by gel electrophoresis and quantified using Quant-iT PicoGreen double-stranded DNA Assay (Life Technologies) using a Fluoroskan Ascent FL fluorometer (Thermo LabSystems).

Double-digest RADseq (Baird et al. 2008) was performed as per methods previously outlined (Elshire et al. 2011) and described in full elsewhere (Poland et al. 2012). Briefly, two restriction enzymes were used (*Pst*I and *Msp*I) to digest genomic DNA. Digested DNA was then ligated with adapters and barcodes for individual identification then amplified by PCR. For the offspring, uniquely barcoded individuals were then combined in equimolar proportions into eight pools, each pool containing 25 individuals. Pools were each sequenced on a single lane on a HiSeq2000 at Génome Québec Innovation Centre (McGill University, Montréal). In order to obtain deeper sequencing of the parents, each parent individual was sequenced using an Ion Torrent at the sequencing platform at IBIS (Institut de Biologie Integrative et des Systemes, Universite Laval, Quebec City). This platform change between F1 and F2 individuals occurred due to equipment availability, but extra precaution was taken to ensure proper correspondence of loci (*see below*).

#### Bioinformatic pipeline and reduced genome *de novo* assembly

Raw reads were inspected for overall quality and presence of adapters with fastqc (http://www.bioinformatics.babraham.ac.uk/projects/fastqc/). Adapters were removed and raw reads were truncated to 80pb using cutadapt v.1.9 Dev.0 (Martin 2011). Reads were demultiplexed, by barcodes, and quality trimmed to 80bp using the stacks v.1.32 (Catchen et al. 2011; 2013) *processradtags* module. The ploidy-informed empirical procedure was used (Ilut et al. 2014) to optimize *de novo* assembly. Sequence similarity was explored to find the optimum clustering threshold, which is highly important for pseudotetraploid salmonid *de novo* assembly (see Additional File S1 for pipeline parameters). Data from each individual were grouped into loci, and polymorphic nucleotide sites were identified with the *ustacks* module. The catalog construction used all loci identified across the parents. Differentially fixed loci (i.e. monomorphic loci among parents) were allowed to merge as a single locus when no mismatches were found *(cstacks)*. Loci from parents and offspring were matched against the parental catalog to determine the allelic state at each locus in each individual in *sstacks*. To improve the quality of the *de novo* assemblies produced in stacks and to reduce the risk of generating nonsensical loci with repetitive sequences and paralogs, we used the correction module *rxstacks*. The log-likelihood threshold for *rxstacks* was chosen based on the distribution of mean and median log-likelihood values. After the correction module, the catalog and individuals' matches were rebuilt with the corrected individuals files. The *genotypes* module of stacks was used to output markers along with their allelic state and raw genotypes. The markers were translated using the function *genotypessummary.R* of stackr (v.0.2.1; Gosselin & Bernatchez, 2015) into fully-or semi-informative markers types, specifically the four types of markers that permitted in the outbreeding design: *ab* x *ac, ab* x *ab, ab* × *aa* and *aa* × *ab* (Wu et al. 2002).

#### Pre-mapping quality control

Several steps of quality control were performed based on recommendations (van Ooijen & Jansen 2013). Pre-mapping quality control consisted of excluding individuals with>30% missing data (22 progeny), monomorphic loci and loci with an incomplete segregation pattern inferred from the parents (i.e. missing alleles) using the *genotypes summary.R* function. This function was also used to filter errors in the phenotype observations of markers with a segregation distortion filter using a chi-square goodness-of-fit test *(filter.GOF)*. With heterozygous parents, not all of the markers contribute equally to the construction of the map, because linkage phases change across loci (van Ooijen & Jansen 2013). Therefore, tolerance for genotyping errors (goodness-of-fit threshold: 12 to 20) and missing genotypes (50% to 90% thresholds) were also explored with *genotypessummary.R*.

#### Linkage mapping and post-mapping quality control

The linkage map was first built in joinmap v4.1; (van Ooijen 2006) using the pseudo-testcross approach (Grattapaglia & Sederoff 1994; van Ooijen & Jansen 2013) that only uses the markers segregating in a uni-parental configuration (i.e. *ab* × *ac* and *ab* × *ab* markers are excluded at this step). Then a consensus map including the bi-parental configuration markers was produced with the joint data analyses as a CP population type (cross pollinator, or full-sib family) using the multipoint maximum likelihood mapping algorithm for marker order (van Ooijen 2011; van Ooijen & Jansen 2013). The pseudo-testcross maps were used for confirmation. Separate maximum likelihood maps were generated for each parent, and only the female map was retained, as is typical for salmonid mapping studies (Kodama et al. 2014). Markers were grouped with the independent LOD option of JOINMAP with a range of 15 to 40 LOD, for the minimum and maximum threshold, respectively. A total of 42 linkage groups (LGs) were defined by evaluating stability of marker numbers over increasing consecutive LOD values. This number of LGs corresponds to the expected chromosome number of Brook Charr (2n=84). During mapping, the stabilization criterion was monitored in the session log with the sum of recombination frequencies of adjacent segments and the mean number of recombination events. Default mapping parameters usually performed well with the smaller LG, but for larger LG the stabilization was not always reached, so more EM cycles and longer chains per cycle were used. For full details of parameters used in JOINMAP, see Additional File S1.

Problematic markers, unlinked markers and small linkage groups were inspected and tested by using several JOINMAP features, including *crosslink*, *genotype probabilities*, *fit* and *stress*. As recommended by van Ooijen and Jansen (2013), detection of errors in ordering and genotyping along with marker exclusion followed these criteria: (i) oversized LG, which can occur with high marker numbers; (ii) incidence of improbable genotypes e.g. double recombinants (Henning et al. 2014), also inspected using the *countXO* function of R/qtl (Broman et al. 2003); (iii) drastic changes of orders; and (iv) low levels of fit or high levels of stress. Maps were inspected for distortion before and after manual exclusion of markers. Mapping distances (cM) were calculated using the Haldane mapping function.

### MapComp

#### Map comparison through intermediate reference genome

In order to compare the Brook Charr map to other salmonid maps, published linkage map datasets were collected (see the Code and Pipeline Availability section), including information on marker name, sequence, linkage group and cM position. Comparisons of linkage group orthology and order were investigated for Chinook Salmon (Brieuc et al. 2014), Pink Salmon (Limborg et al. 2014), Chum Salmon (Waples et al. 2016), Rainbow Trout (Miller et al. 2012; Palti et al. 2015), Sockeye Salmon (Everett et al. 2012; Limborg et al. 2015), Coho Salmon (Kodama et al. 2014), Atlantic Salmon (Lien et al. 2011), Lake Whitefish (Gagnaire et al. 2013), and the salmonid WGD sister outgroup Northern Pike (Rondeau et al. 2014) (Table 1).

The basic workflow of MapComp is shown in Figure 1. First, all marker sequences were combined into a single fasta file and mapped to a reference genome, here the Rainbow Trout scaffolds (http://www.genoscope.cns.fr/trout/data/ added 28-Apr-2014) (Berthelot et al. 2014) and the Atlantic Salmon assembly ICSASG_v2 (AGKD00000000.4; Lien et al. 2016) using *BWA mem* (Li & Durbin 2009). Matches to the reference were only retained when mapping quality score (MAPQ) was≥10 and a single match was found in the target genome. When two markers (i.e. one from each species) mapped to the same reference genome scaffold or contig, the two closest markers were taken as a marker pair. Markers were paired without replacement (i.e. once the closest marker pair was selected, other markers also pairing with the marker that has now been paired were then discarded). Each marker pair was then added to an Oxford grid. The pipeline developed for MapComp is available at https://github.com/enormandeau/mapcomp/.

**Table 1.**
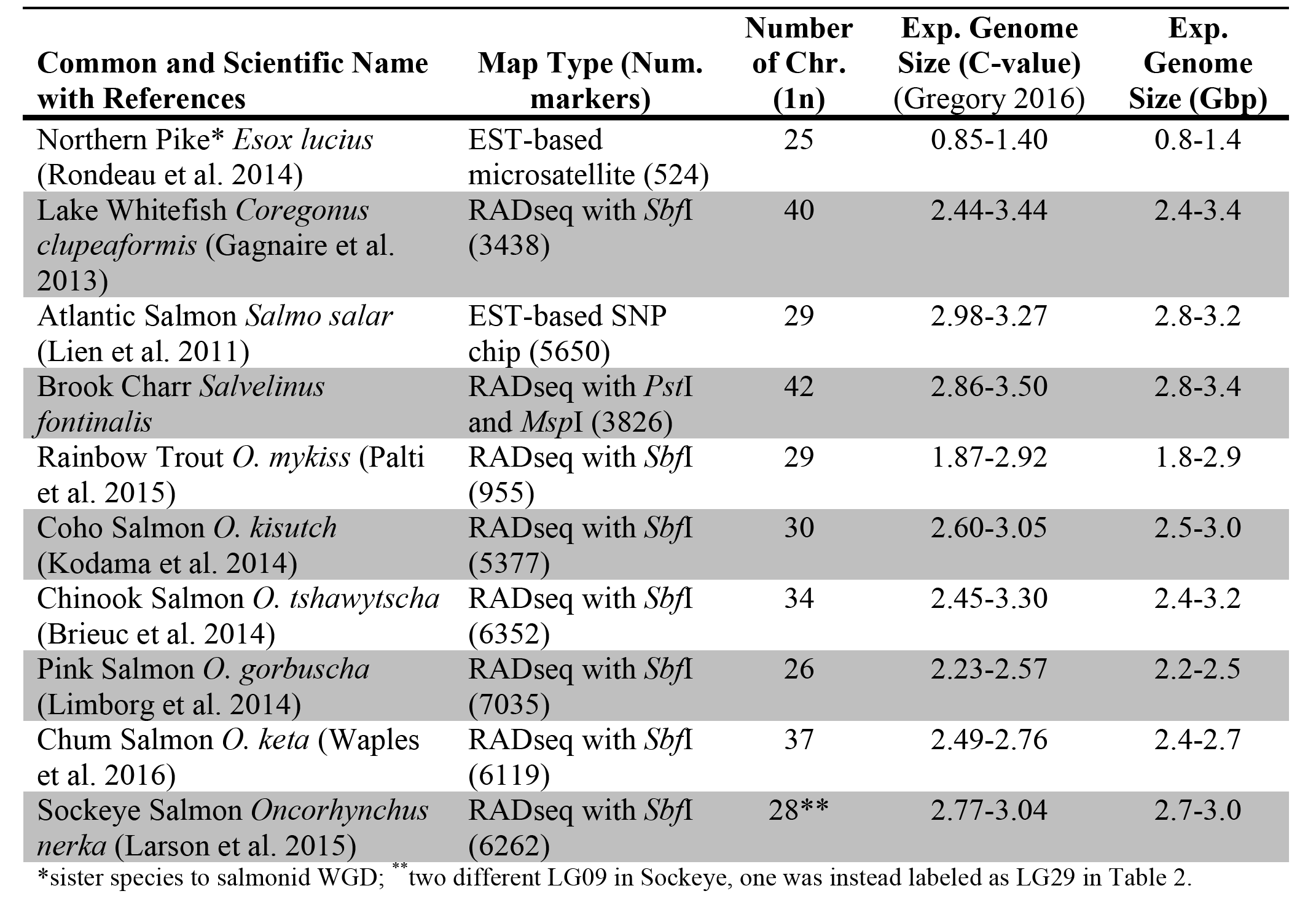
**Overview of compared species**. The common and scientific name for each species in the analysis are displayed along with the source of the genetic map, the type of map and number of markers, the chromosome number for the species, and expected genome size (C-value and Gbp) obtained from (Gregory 2016).

To identify homeology relationships, where two chromosome arms originate from the same pre-duplicated chromosome arm, comparisons between Northern Pike *E. lucius* and the other salmonids were conducted. This required changing some parameters in MapComp to allow for multiple hits from the non-duplicated Northern Pike map against the Atlantic Salmon and Rainbow Trout reference genome intermediates, as each marker could be present in at least duplicate in the salmonid genome. Specifically, the mapping quality threshold was lowered (MAPQ≥2) and mapping against more than one locus in the reference genome was permitted.

**Fig. 1.**
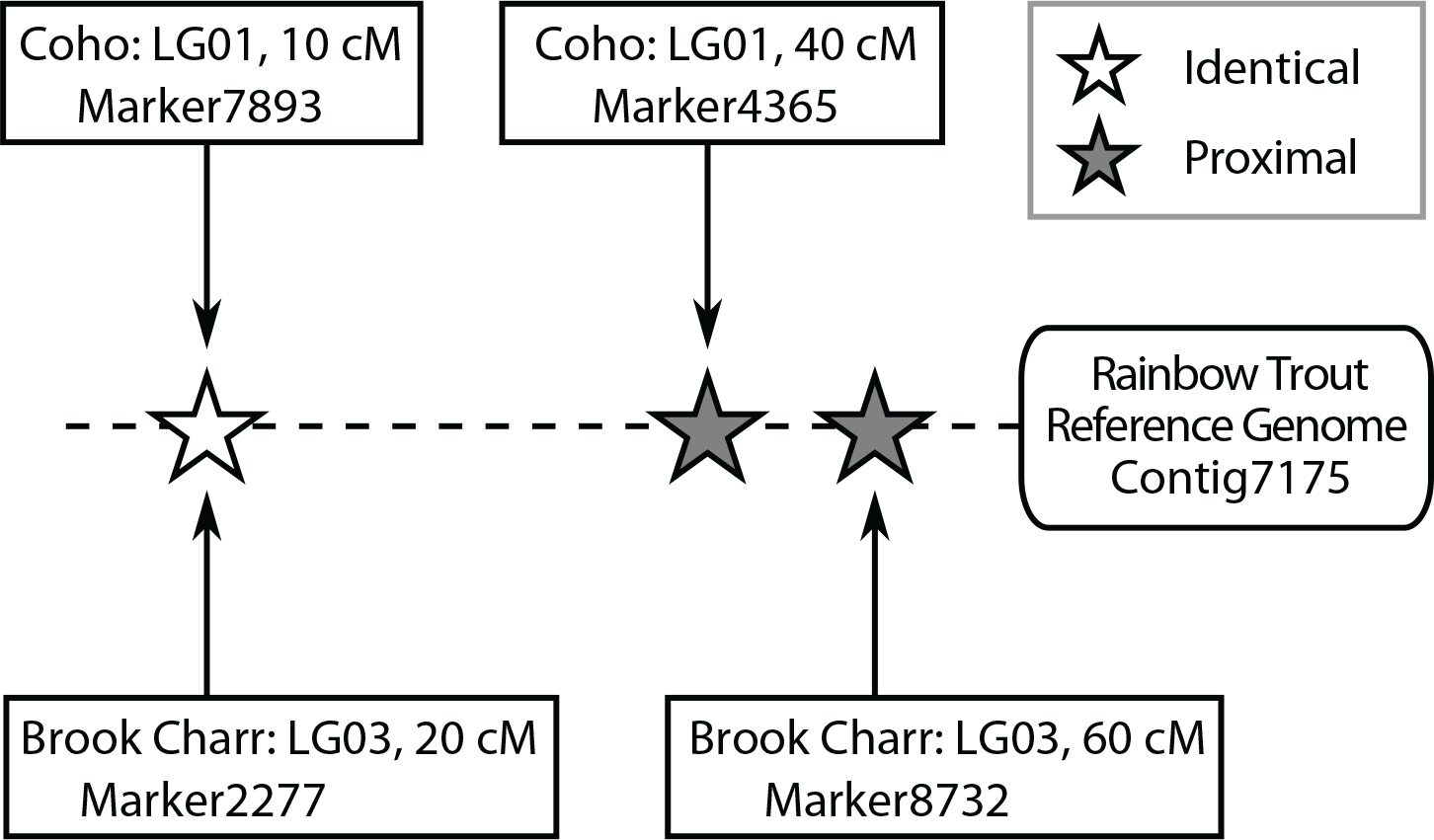
**Schematic of MapComp using a reference genome to pair markers**. MAPCOMP compares genetic maps from two different species by mapping marker sequences against a reference genome, then retaining high quality mappings that only hit against one locus in the genome. Markers from each species are paired if they hit against the same contig/scaffold by taking the closest two markers together as each pair. Each marker is paired without replacement, and so any other marker that was second closest to the now-paired marker is discarded. This method captures identical markers (white star in image) and non-identical markers (grey stars). Finally, the linkage group and cM position of each marker is plotted in an Oxford grid. Note that the marker names and contig ID in the schematic are for demonstration purposes only and do not reflect actual pairings.

#### Characterization of orthology and homeology between chromosome arms

Orthology of chromosome arms between Chinook Salmon and Coho Salmon maps identified previously using homologous markers (Kodama et al. 2014) was confirmed using MapComp. Chinook Salmon and Coho Salmon were then individually compared with Brook Charr to identify corresponding chromosome arms in Brook Charr. Once these orthology relationships were obtained, the Brook Charr map was compared with Pink Salmon, Sockeye Salmon, Chum Salmon, Rainbow Trout and Atlantic Salmon. Orthology was identified in Lake Whitefish using a consensus approach, where results from comparisons of Lake Whitefish with multiple different species were considered for unambiguous determination of orthology. Homeologs were also identified using a consensus approach, and the original Northern Pike linkage groups were given .1 or .2 designations to represent the duplicated chromosomes. These results were compared to the results of Rondeau et al. (2014), in which BLAST was used with Atlantic Salmon linkage groups against the Northern Pike genome to identify salmonid WGD homeologs.

#### Identification of putative inversions

Plots from MapComp were visually inspected for inversions. During linkage mapping, when markers do not fit in the linkage group, they can be placed at the distal ends of the linkage group (Henning et al. 2014). Therefore, to avoid the erroneous identification of inversions, evidence for inversions was only considered when non-inverted regions flanked the inverted region. As the analysis is based on linkage maps and not assembled genomes, all inversions were considered putative. Furthermore, phylogenetic relationships and inversion conservation across species were considered (i.e. when an inversion was identified within multiple species within a lineage). Centromere locations were obtained from Chinook Salmon (Brieuc et al. 2014) to allow the characterization of selected inversions as either pericentric (involving the centromere) or paracentric (not involving the centromere).

#### Conservation of rearrangements and identification of full coverage of linkage groups

The conservation of chromosomal rearrangements among the salmonids was analyzed by using the most taxonomically complete phylogeny of the salmonids (Crête-Lafrenière et al. 2012) but with Rainbow Trout as the outgroup to the other *Oncorhynchus* clades as reported previously (Kinnison & Hendry 2004), and with the still-debated clade containing Pink Salmon, Chum Salmon and Sockeye Salmon arranged in the most parsimonious phylogeny in terms of the numbers of required fusions/fissions. The analysis of metacentric conservation was based on the analysis of conservation in Coho, Chinook, Rainbow Trout and Atlantic Salmon (Kodama et al. 2014), but re-analyzed using MapComp and additional maps in the present study (i.e. Pink Salmon, Chum Salmon, Sockeye Salmon, Brook Charr and Lake Whitefish). To confirm that a metacentric chromosome was completely present, for conserved metacentric identification we required evidence from both sides of the centromere.

## Results

### Generation of a Brook Charr linkage map

On average, 10M single-end reads were obtained for each parent and 5M for each individual offspring. Using STACKS v1.32 (Catchen et al. 2011), 6264 segregating markers were identified, each containing one to five SNPs. Missing data per marker followed a heavy-tailed distribution, having a mode of 10 individuals genotypes missing for ~700 markers. Female, male and consensus genetic maps were generated, but the female-specific map (n=3826 markers) was retained as the final map, as is typical for salmonids due to low recombination rate (e.g. Kodama et al. 2014; Brieuc et al. 2014) and increased residual tetrasomy in males (Allendorf et al. 2015).

A total of 42 linkage groups were characterized in the female map (Figure 2), corresponding to the expected haploid chromosome number for Brook Charr (Phillips & Ráb 2001). On average, metacentric linkage groups were 270 cM (range=185–342 cM) containing 126 markers (range=107–175 markers), whereas acrocentric linkage groups were 156 cM (range=65–230 cM) containing 83 markers (range=33–134). The total length of the female map was 7453.9 cM. Descriptive statistics for the linkage groups are in Additional File S2. This size is in the range of other high-density salmonid maps, such as the Coho Salmon linkage map (6596.7 cM) (Kodama et al. 2014), although is larger than the Chinook Salmon map (4164 cM) (Brieuc et al. 2014). The female map contains 3826 markers with the following marker types, as defined by Wu et al. (2002): 254 fully-informative *(ab* × *ac)*, 954 semi-informative *(ab* x *ab)* and 2618 fully informative in female parent *(ab* x *aa)*. The female map is in Additional File S3. The consensus map contained an additional 2385 markers that were informative in the male parent (*aa* × *ab*) but although these markers were in the correct linkage groups, they did not position well within the linkage group. This is most likely due to the low recombination rate known to occur in male salmonids, as almost complete crossover interference can occur within male salmonids during meiosis (Naish et al. 2013).

**Fig. 2.**
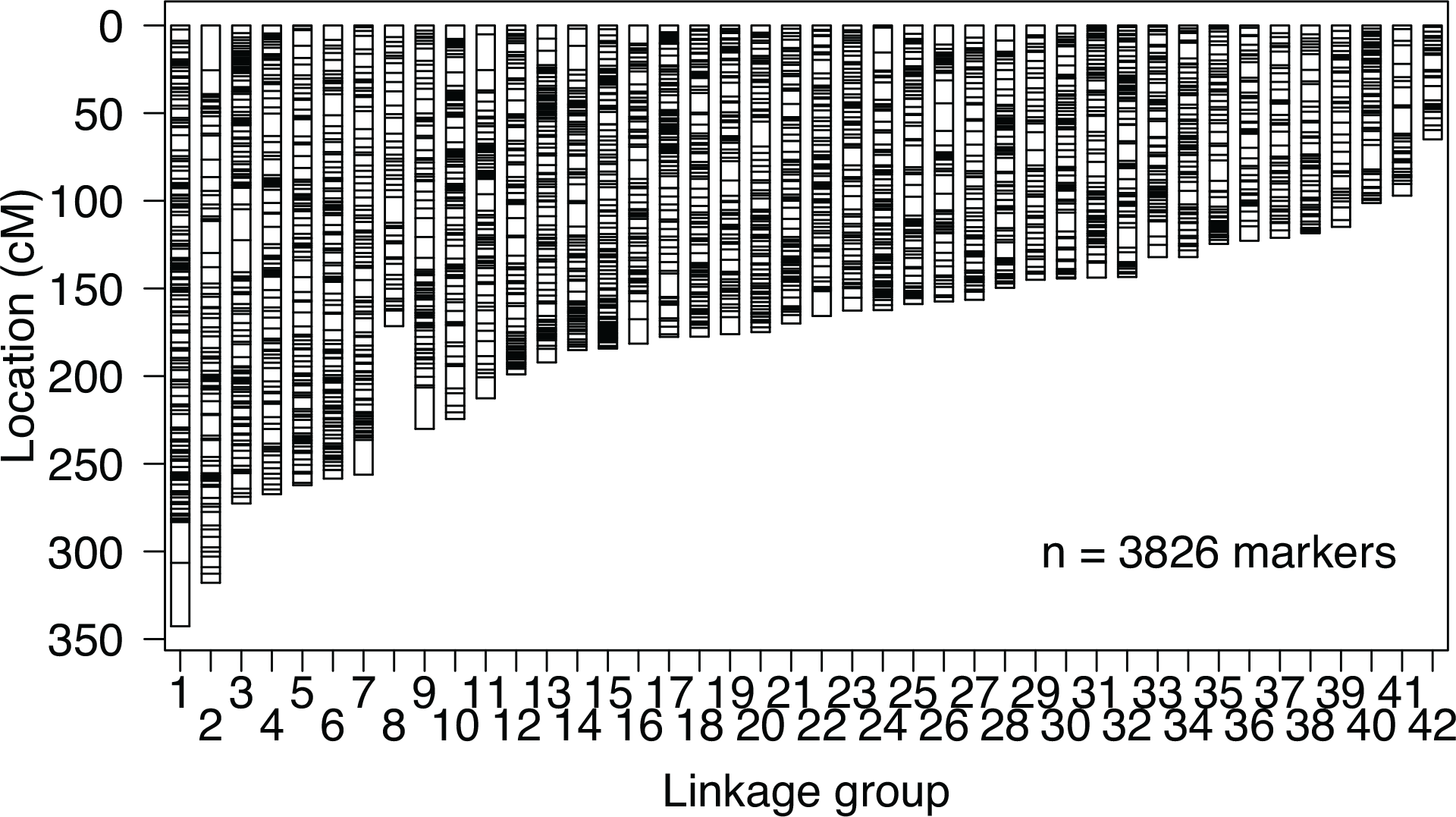
**Brook Charr *Salvelinus fontinalis* linkage map**. Eight metacentric (LG1–8) and 34 acrocentric linkage groups (LG9–42) were identified in the female map. Horizontal lines within each linkage group are markers (total=3826 markers).

### Identification of orthologous chromosome arms among the salmonids

Assignment of linkage groups to chromosome arms has been performed using fluorescence *in situ* hybridization with BAC probes for Atlantic Salmon (Phillips et al. 2009), and orthology has been designated using homologous microsatellite and RADseq markers (using the same library preparation protocols) among Chinook Salmon, Coho Salmon, Rainbow Trout and Atlantic Salmon (Danzmann et al. 2008; Phillips et al. 2009; Naish et al. 2013; Kodama et al. 2014; Brieuc et al. 2014) and recently Sockeye Salmon (Larson et al. 2015). A full comparison across all existing maps has yet to be completed. The low-density linkage map of the Northern Pike *E. lucius* has been compared with Atlantic Salmon (Rondeau et al. 2014), but not yet with the rest of the salmonids. Details on the linkage maps and species used in this analysis, including expected genome sizes (Gregory 2016) are provided in Table 1.

To begin orthology designation of linkage groups, Chinook Salmon and Coho Salmon linkage maps were used to compare with the map of Brook Charr using MapComp pairing markers through the Rainbow Trout genome (Berthelot et al. 2014) (see MapComp schematic in Figure 1, and Methods for full details). All chromosome arms (NF=50) were identified unambiguously in Brook Charr (Figure 3; Table 2). The Brook Charr linkage map was then compared with linkage maps of Sockeye Salmon, Chum Salmon, Pink Salmon, Rainbow Trout and Atlantic Salmon (Table 2). In a few rare cases where orthology with Brook Charr was not obvious, species were also compared to Chinook Salmon or others to clearly indicate the corresponding chromosome arm. One chromosome arm in Rainbow Trout required the use of a second Rainbow Trout linkage map to unambiguously identify orthology (see Table 2; Miller et al. 2012). Most arms were also identified in the more distantly related Lake Whitefish, but seven arms remained unidentifiable.

**Table 2.**
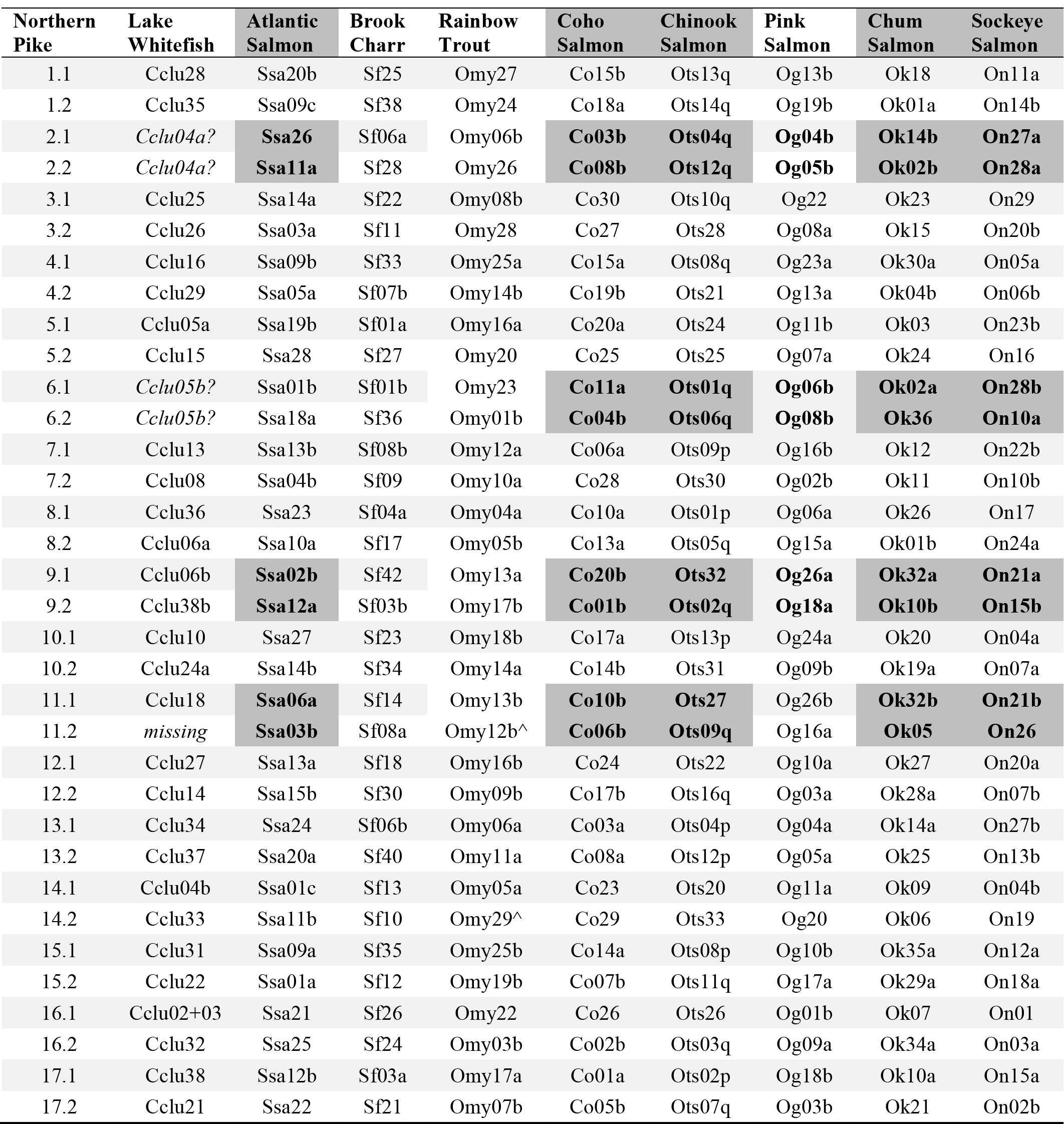
**Orthologous chromosome arms across the salmonids**. All orthologous relationships among salmonids and the pre-duplicated Northern Pike are displayed as identified by MapComp. Bold/grey-shaded species are those with studies that specifically tested for residual tetrasomy using duplicate markers or sequence similarity, and within these species, the bold/grey-shaded homeologs are those with evidence of residual tetrasomy in the original studies (see Table 1 for references). Note that Northern Pike only has 25 chromosomes, but here each chromosome is listed twice to accommodate the duplicate orthologs in the other species.

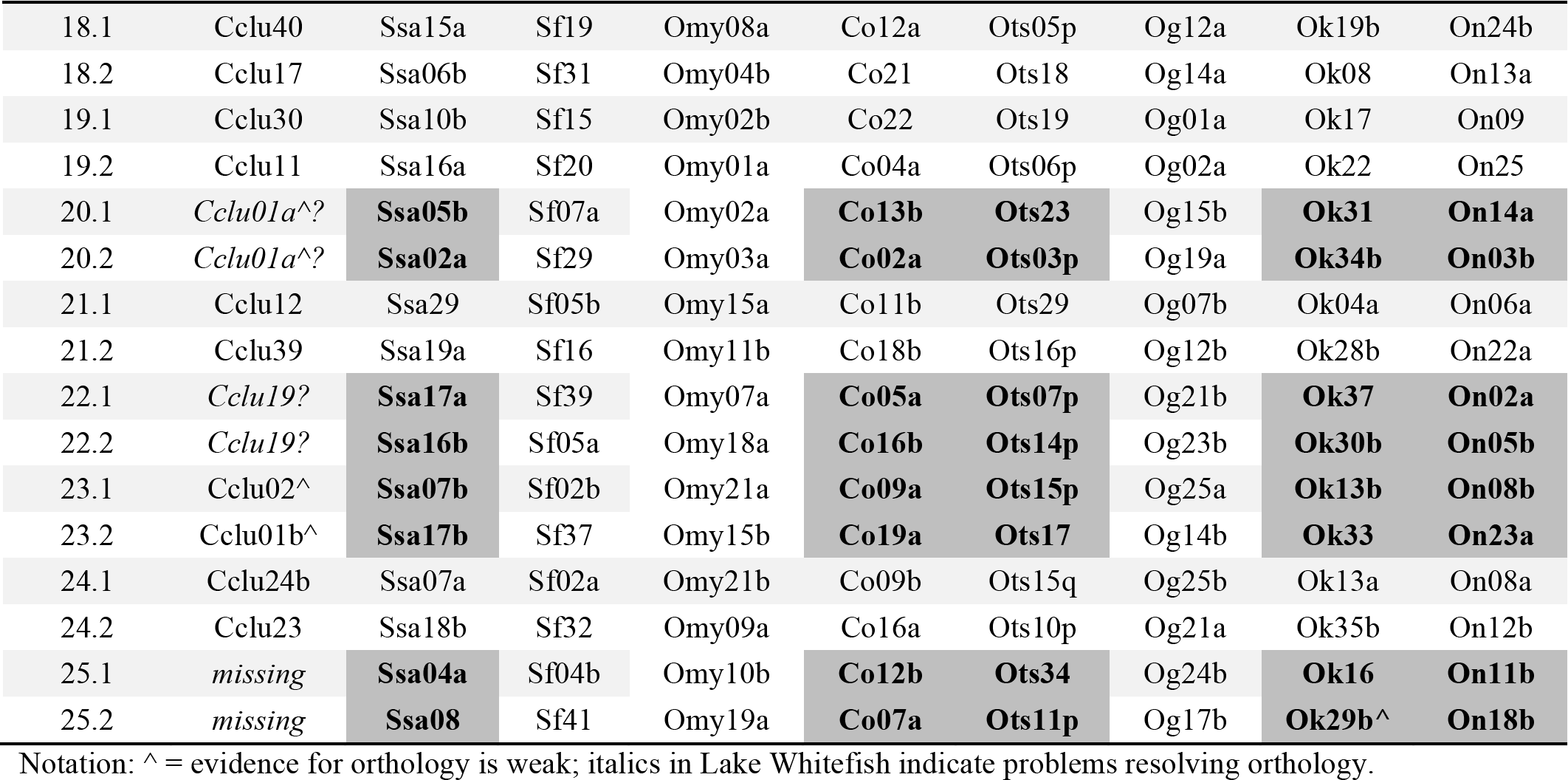

In Brook Charr, a total of eight metacentric and 34 acrocentric chromosomes were expected from salmonid cytogenetics (Phillips & Ráb 2001; Lee & Wright 1981) and all were identified here (Table 2), increasing the resolution of the Brook Charr linkage maps from existing microsatellite and low density SNP-based linkage maps (Sauvage et al. 2012a; Timusk et al. 2011). Since Brook Charr has the fewest metacentric chromosomes of the species characterized here, often two acrocentric chromosomes in Brook Charr correspond to two fused arms on the same metacentric chromosome in another species. In some cases, due to tandem chromosome fusions observed in Atlantic Salmon (Phillips et al. 2009) three linkage groups in Brook Charr correspond to one linkage group in Atlantic Salmon. For example, Brook Charr Sf33, Sf35, Sf38 are in tandem fusions in Atlantic Salmon Ssa09. We compared orthology identified with MapComp between Coho Salmon and Chinook Salmon with an analyses that used homologous markers (Kodama et al. 2014), and found the same correspondence. We identified discordant results for five putative orthologous chromosomes between Chinook Salmon and Atlantic Salmon, and for two putative orthologous chromosomes between Chinook and Rainbow Trout that were based on earlier studies. The rest of the results among these species corresponded between the studies (total=50 orthologous relationships in four species). Additionally, orthology was determined for Pink Salmon, Chum Salmon, Sockeye Salmon and Lake Whitefish.

**Fig. 3.**
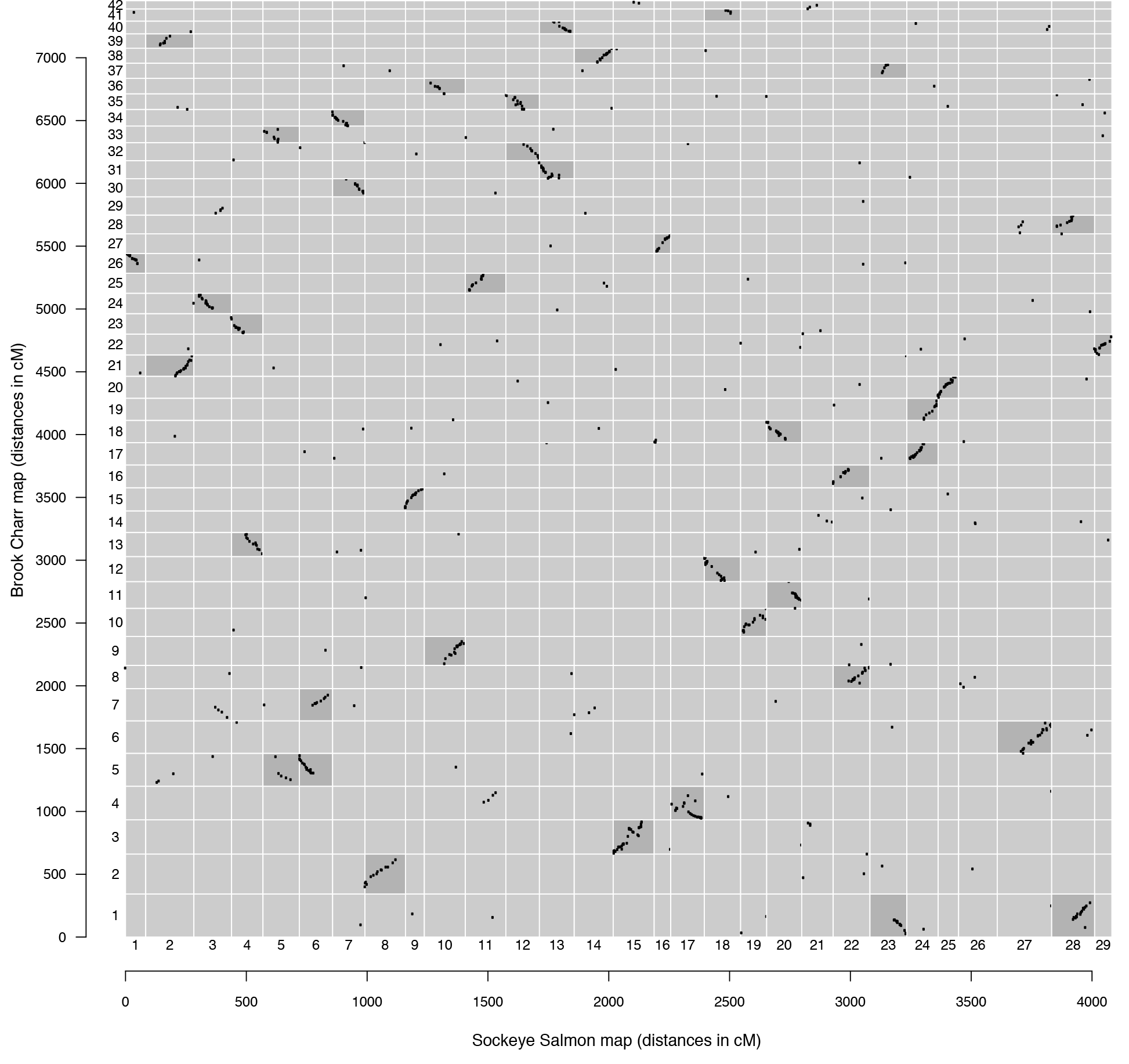
**MapComp determination of orthologous chromosome arms**. Brook Charr compared with Sockeye Salmon with markers paired through the Rainbow Trout genome identifies orthology between chromosome arms. A putative inversion can be seen between Brook Charr LG03 and Sockeye Salmon LG15.

### Homeologous chromosome identification

To identify homeologous chromosomes (i.e. chromosome arms originating from the same preduplicated chromosome), the genetic map of Northern Pike was compared with the maps of all species using MapComp as described in the Methods. All homeologous pairs in Atlantic Salmon identified by MapComp using the Rainbow Trout intermediate reference genome were concurrent with those originally identified (Rondeau et al. 2014), but here were also extended to all other species (Table 2).

Homeologous chromosomes that undergo residual tetrasomy can be identified by mapping isoloci (i.e. markers recombining between homeologous chromosomes) in haploid crosses, as has been done for Atlantic Salmon, Coho Salmon, Chinook Salmon, Sockeye Salmon and Chum Salmon (see Table 1 for references). Sequence similarity between homeologous chromosomes has also been used to identify residual tetrasomic pairs in the analysis of the Atlantic Salmon genome (Lien et al. 2016). As can be seen from the orthologous chromosomes in Table 2, the same eight pairs of homeologous chromosomes are residually tetrasomic in the *Oncorhynchus* species, and seven of these are still residually tetrasomic in Atlantic Salmon. Without a haploid cross for our Brook Charr map, here we cannot specify whether any homeologs still undergo residual tetrasomy in this species. The homeologs identified using Northern Pike with MapComp were compared with those identified using isoloci in mapping crosses. All eight pairs of homeologs identified in Chinook Salmon (Brieuc et al. 2014) were confirmed as originating from the same ancestral chromosome. The eight pairs of homeologs identified in Coho Salmon (Kodama et al. 2014) and Sockeye Salmon (Larson et al. 2015) were also confirmed here. In addition, all other homeologies (total=25 pairs) in all evaluated species were also identified using MapComp, with the exception of the aforementioned unidentifiable chromosome arms of Lake Whitefish (Table 2). Interestingly, all of the missing orthology designations in Lake Whitefish are, without exception, those with residual tetrasomy in *Oncorhynchus* (Table 2).

### Conserved and species-specific chromosome rearrangements

Shared rearrangements among species in a clade (e.g. fusion events) are likely to have occurred prior to the diversification of the clade, as demonstrated for nine metacentric fusions in Coho Salmon and Chinook Salmon (Kodama et al. 2014). Here, chromosome arm orthology designation furthered this analysis, allowing the inclusion of Brook Charr and Lake Whitefish as well as the clade containing Sockeye Salmon, Pink Salmon and Chum Salmon within *Oncorhynchus*. We identified 16 different fusion events conserved in at least two species, five fission events conserved in at least two species, 87 species-specific fusion events, and five species-specific fission events (Figure 4). For clarity, when discussing fusions and fissions here we use chromosome names from the Northern Pike chromosomes to refer to chromosome arms, including the duplicate designation (e.g. 1.1/1.2 or 3.1/3.2), as shown in Table 2. The phylogeny in Figure 4 was adapted from previous literature (Stearley & Smith 1993; Kinnison & Hendry 2004; Crête-Lafrenière et al. 2012). For the clade containing Pink, Sockeye and Chum Salmon, in which the sister relationships remain unclear (Kinnison & Hendry 2004), we present the most parsimonious phylogenetic relationship in terms of required number of fusion/fission events. With Pink Salmon as the sister species to the Chum and Sockeye Salmon clade, this requires three fewer fission or fusion events, and one fewer fission of a conserved metacentric chromosome, the 7.1–11.2 fusion.

**Fig. 4.**
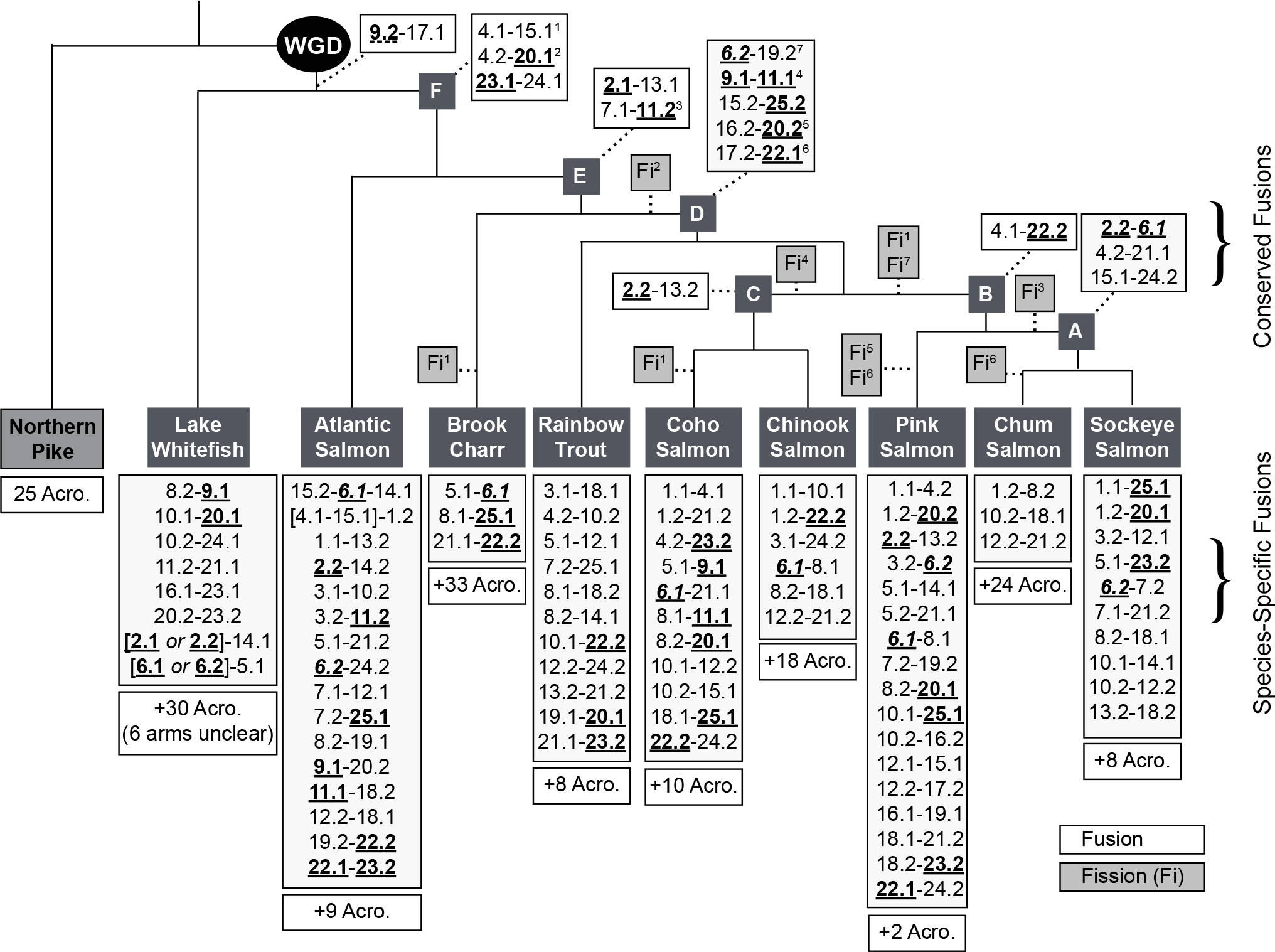
**Fusions and fissions across the salmonid lineage**. Different fusions and fissions have occurred during the evolution of the salmonids. White boxes display the fusion events, where the orthologous chromosomes for all species are named according to the corresponding Northern Pike linkage group ID, with .1 or .2 to correspond with the post-duplicated salmonid chromosomes. Bold and underlined chromosome numbers are the homeologous pairs that still undergo residual tetrasomy, and the italicized chromosome (6.1/6.2) is not residually tetrasomic in Atlantic Salmon but is in the Pacific salmonids. Above the species names are conserved fusions, whereas below are the species-specific fusions. Also shown are fissions in light grey with the notation (Fi^x^), where *x* corresponds to the superscript in the original fusion (e.g. 4.2–20.1 at point (F) in the phylogeny is divided at Fi^2^ prior to point (D)). The phylogeny is adapted from (Stearley & Smith 1993; Kinnison & Hendry 2004; Crête-Lafrenière et al. 2012), with minor modifications to the relationships within the Pink, Chum and Sockeye Salmon clade, as described in the results. Branch lengths are not to scale and are for illustrative purposes of relationships between species only.

The oldest identified rearrangement is the 9.2–17.1 fusion event that is conserved in all species investigated (see Figure 4). Conserved in all species except Lake Whitefish is the 23.1–24.1 fusion (see F in Figure 4). Another metacentric fusion event at this same point in the phylogeny was also identified (4.2–20.1) that is still present in both Atlantic Salmon and Brook Charr, but not in any members of *Oncorhynchus*, suggesting that the a fission occurred prior to the speciation of any members of *Oncorhynchus* (Fi^2^ in Figure 4). One fusion (2.1–13.1) is present in Brook Charr and all *Oncorhynchus* spp. (fused at E in Figure 4). Another fusion at this same point (7.1–11.2; E in Figure 4) is found in all descendants except for the Chum and Sockeye Salmon clade (fission Fi^3^ in Figure 4).

More recent rearrangements include five fusions prior to the speciation of the *Oncorhynchus* clade (D in Figure 4). One is present in all *Oncorhynchus* species (15.2–25.2), another is present in all *Oncorhynchus* species except Pink Salmon (16.2–20.2; see Fi^5^ in Figure 4), another underwent fission in the Chinook/Coho lineage (9.1–11.1; see Fi^4^) and one underwent fission in Pink and Chum Salmon (17.2–22.1; see Fi^6^). Conserved fusions were found within the *Oncorhynchus* lineage as well, including one fusion in the Chinook/Coho lineage, (2.2–13.2; C in Figure 4) and three fusions prior in the Sockeye/Chum lineage (A in Figure 4). Each species also has had species-specific fusions, ranging in number from only three fusions in Chum Salmon and three in Brook Charr to up to 17 in Pink Salmon and 18 in Atlantic Salmon (Figure 4).

Some rearrangements are more complex and thus it is more difficult to unambiguously describe their history. For example, a three chromosome arm fusion in Atlantic Salmon occurred through a single fusion (4.1–15.1) that either fused once prior to the divergence of Atlantic Salmon and underwent three different fission events (Fi^1^ in Figure 4), or fused three independent times with the same fusion partner. It is not clear which of these possibilities is correct, but in Figure 4 we display the first and more parsimonious scenario. In Atlantic Salmon, after the metacentric fusion, an additional fusion occurred, adding a third chromosome arm (1.2 with [15.1–4.1]). Three other different fusions appeared to have occurred two independent times: 8.2–18.1 in Chinook and Sockeye; 12.2–21.2 in Chinook and Chum; and 7.2–25.1 in Atlantic Salmon and Rainbow Trout. For each of these multiple independent origins, the alternate explanations are possible but less parsimonious. Although these few independent origin cases are not entirely clear, we display the most parsimonious rearrangements, requiring the fewest independent fusions/fissions in Figure 4.

### Putative lineage-specific inversions

Several inversions flanked by non-inverted regions were revealed between linkage maps, suggesting the presence of chromosomal segment inversions (Figure 5). These putative inversions are more supported when phylogenetically conserved. Future genome assemblies for the species involved will be valuable for further inversion identification. A striking putative inversion was identified in one of the metacentric chromosomes conserved across all evaluated salmonids (9.2–17.1; Figure 4). An inversion near the center of the linkage group is present in only the Pink, Chum and Sockeye Salmon lineage. This inversion is visible in the Oxford grids between these species and Coho, Chinook Salmon and Brook Charr (see Figure 5a–b). As a result, the conformation observed in Pink, Chum and Sockeye Salmon is likely the derived form. Rainbow Trout does not indicate the inversion against the ancestral conformation, but also does not indicate the inversion with Pink, Chum and Sockeye Salmon, as there is a gap with no marker pairs available at the inverted locus in the Rainbow Trout linkage group.

**Fig. 5.**
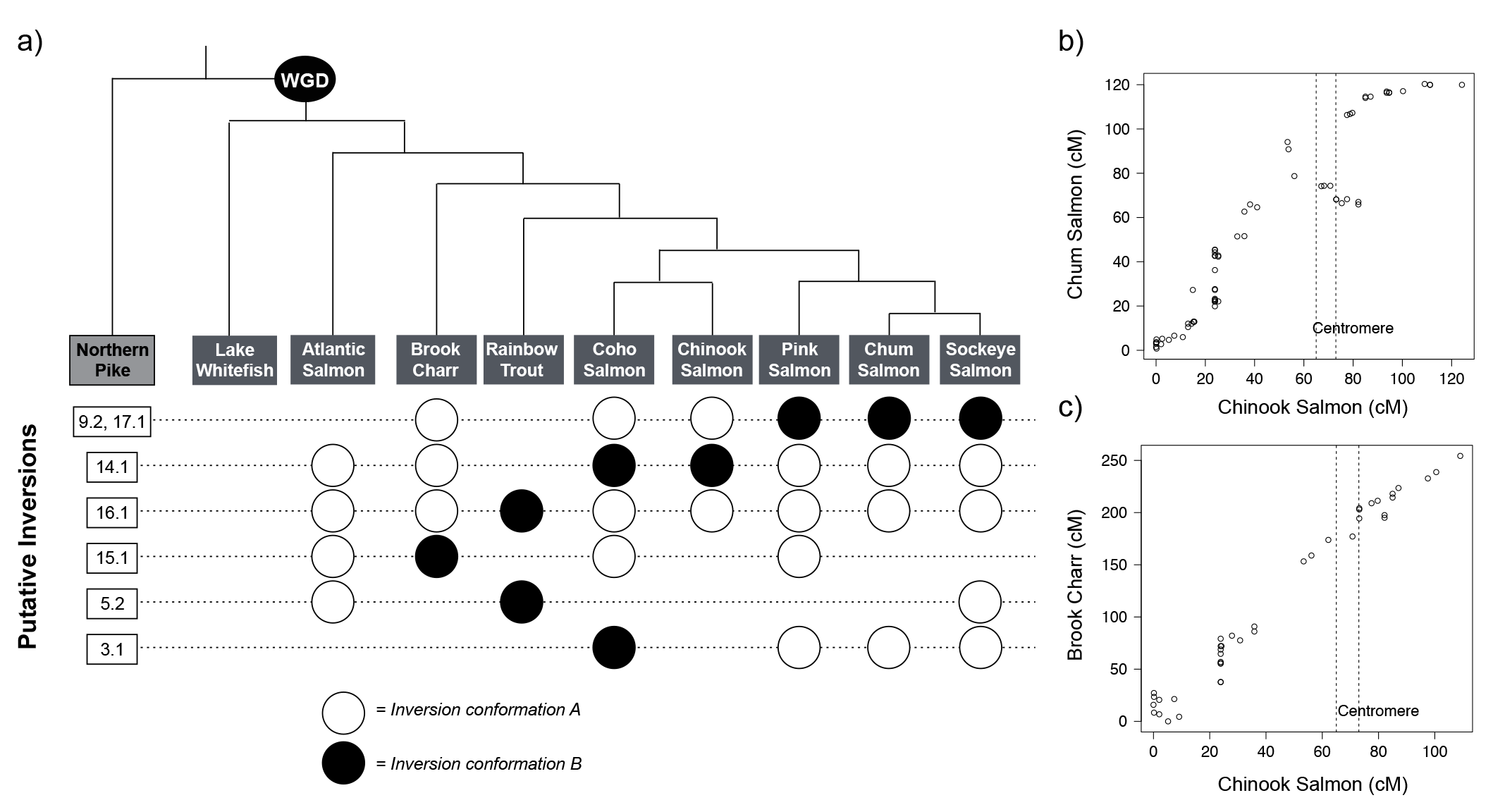
**Putative conserved and species-specific inversions**. a) The salmonid phylogeny is shown highlighting six different inversion events, each listed according to the Northern Pike chromosome and represented on a dotted line below the phylogeny. Per line (inversion event), white circles indicate the more common or likely ancestral inversion conformation, and black circles the less common and more likely derived inversion conformation. If no circle is present, the inversion was not visible in the linkage map of that species. The putative pericentric inversion across the fusion 9.2–17.1 is displayed in (b) showing two species (Chum and Chinook Salmon) with different inversion conformations, and in (c) for two species with the same conformation (Brook Charr and Chinook Salmon). Predicted centromere positions previously identified in Chinook Salmon (Brieuc et al. 2014) are also shown in (b-c). Full names for species are defined in Table 1, the phylogeny is as described in the Results and Figure 4, and probable genes within the 9.2–17.1 inversion are shown in Additional file S4.

To further characterize the 9.2–17.1 inversion, centromere locations obtained from Chinook Salmon (Brieuc et al. 2014) were compared to the location of the inversion. The inverted region corresponds to the Chinook Salmon linkage group 0ts02 between ~49–82 cM (see Figure 5b) and the centromere for this linkage group was estimated to be between 65–73 cM. This inversion therefore most likely contains the centromere (i.e. a pericentric inversion). This inverted region between Chinook Salmon and Chum Salmon was visible when using either the Rainbow Trout or Atlantic Salmon genome as the intermediate reference genome, and had evidence from 12 markers mapping through two different scaffolds. To identify the genes that may be contained within this inverted region, the mapped locations on the Atlantic Salmon genome of the markers at the distal ends of the inverted region were taken. As the Atlantic Salmon genome has been annotated (Lien et al. 2016), this region of the genome (Ssa12 between 37,048,324–44,754,074 bp) was inspected for gene content. This region (~7.7 Mb) putatively contains 11 genes (based on alignment evidence, here we do not include predicted genes), including *cytokine-like protein 1, solute carrier family 2, facilitated glucose transporter member 9* and *cd8 beta*, among others (see Additional File S4). The exact genes found in this region in the species with the derived conformation of the inversion will require more genomes to be available before exploring further, including the actual breakpoints of the inversion and whether these occur within coding genes. In summary, 9.2 fused with 17.1 in the ancestor of all salmonids investigated here, then an inversion of a segment ~7.7 Mb containing coding sequences occurred across the centromere specifically in the Pink, Chum and Sockeye Salmon lineage. Other inversions were also visible (Figure 5a; Additional File S5). More information on these and potentially new inversions will be obtained as more assembled genomes become available.

### Benefits of *MapComp* versus direct marker comparison and effect of intermediate reference genome

Linkage group orthology between species are typically identified by finding homologous markers using reciprocal best-hit BLAST (Kodama et al. 2014). The method implemented in MapComp, where we accept both identical and proximal markers, leads to a far greater number of retained marker pairs (on average 5-fold; Table 3). For example, between Brook Charr and Chinook Salmon, 907 marker pairs were identified using MapComp, whereas direct mapping identified 190 pairs.

MapComp was tested on both the Rainbow Trout and the Atlantic Salmon genome as intermediate references for pairing markers between maps (Table 3). Results using either genome were highly concordant, having only minor differences in the number of markers mapped and paired. Although the Rainbow Trout genome as an intermediate provided slightly more mapped markers (on average 1.2-fold more than Atlantic Salmon), the Atlantic Salmon genome provided more marker pairs (on average 1.4-fold). This could be due to slight differences in contiguity of the two genomes. As the Atlantic Salmon linkage map sequence information was obtained in EST format, the number of markers mapping from the map to the genome was lower than expected relative to the shorter reads, probably due to the longer sequences and because BWA mem is not a splice-aware aligner.

**Table 3.** **MapComp results using two different intermediate genomes and comparison with results from reciprocal BLAST**. The number and percentage of markers from each species that map to each genome are shown, along with the number of markers pairs between each species and Brook Charr identified by MapComp. Also shown is the number of homologous markers that would have been found between each species and Brook Charr with a reciprocal BLAST approach. Numbers of mappings and pairs were similar when tested on the Rainbow Trout or Atlantic Salmon genome assemblies. *N*/*A* values are present as Brook Charr is not paired against itself.

| Species | Total Markers | Map to Rainbow Trout genome: No. (%) | No. Marker Pairs | Map to Atlantic Salmon genome: No. (%) | No. Marker Pairs | Recip. best-hit BLAST |
| --- | --- | --- | --- | --- | --- | --- |
| Lake Whitefish | 3438 | 1156 (33%) | 346 | 1185 (34%) | 609 | 111 |
| Atlantic Salmon | 5650* | 2776 (49%) | 619 | 3434 (60%) | 1041 | 208 |
| Brook Charr | 3826 | 2321 (60%) | N/A | 2454 (64%) | N/A | N/A |
| Rainbow Trout | 955 | 837 (87%) | 300 | 626 (65%) | 411 | 30 |
| Coho Salmon | 5377 | 3873 (72%) | 813 | 2856 (53%) | 1068 | 182 |
| Chinook Salmon | 6352 | 4663 (73%) | 907 | 3472 (54%) | 1162 | 190 |
| Pink Salmon | 7035 | 4544 (65%) | 841 | 3481 (49%) | 1129 | 210 |
| Chum Salmon | 6119 | 4150 (67%) | 795 | 3139 (51%) | 1049 | 205 |
| Sockeye Salmon | 6262 | 4034 (64%) | 771 | 3138 (50%) | 1061 | 209 |

\*from EST sequences

MapComp parameters can be adjusted for the maximum distance allowed between paired markers. Here we used a maximum distance of 10 Mbp, but most paired markers were at a much smaller distance than this (see marker distance distribution examples in Additional File S6). With a greater phylogenetic distance between species, fewer identical markers were found. For example, the number of identical markers for Chinook and Coho Salmon are high, but the Chinook Salmon comparison with Brook Charr depended more on non-identical markers, as did the comparison between Lake Whitefish and Brook Charr. These parameters can be easily tested by the user, allowing the identification of the optimal settings for individual datasets to permit the greatest number of comparisons without increasing noise in the Oxford grids. Here we found that 10 Mbp provided more markers without a substantial increase in noise.

## Discussion

Linkage maps have many applications, including QTL analysis, assisting genome assembly and comparative genomics. With advances in sequencing technology and techniques (Baird et al. 2008), high quality and dense linkage maps are increasingly available for many species, including non-model species. Dense linkage maps are highly useful for anchoring genome scaffolds to chromosomes (Ming & Man Wai 2015) or for comparative genomics, allowing for information transfer from model to related non-model organisms (Naish et al. 2013). They are also useful for cross-species QTL comparisons (Larson et al. 2015) to understand genome function and evolution, such as that after a whole genome duplication (Kodama et al. 2014).

Salmonids are a valuable taxon for studying genome duplication. Recently, Kodama et al. (2014) characterized several chromosomal fusions and positioned them in the salmonid phylogeny based on the conservation of the fusion across the investigated lineages. This has indicated that structural rearrangements have occurred throughout the evolution of the salmonids, with rearrangements retained from different points in evolutionary history. Here, we further demonstrate the diversity of these rearrangements by identifying all orthologous arms in additional species and genera, and evaluate the most likely timings of rearrangements throughout salmonid evolutionary history.

Residual tetrasomy, or recombination between homeologous chromosomes, occurs through tetravalent formation at meiosis and probably requires that at least one of the homeologous chromosome arms involved is in a metacentric fusion (Wright et al. 1983; Allendorf et al. 2015). As evidence of this, fewer tetravalents form at meiosis when fewer metacentrics chromosomes are present in a species. For example, fewer tetravalents have been observed in comparisons between Rainbow Trout (1n=20 metacentrics) and Brown Trout *Salmo trutta* (1n=10 metacentrics) (May & Delany 2015). This is important to consider for salmonid genome assembly. If a homeologous chromosome pair that is known to be residually tetrasomic in other salmonid species (see Table 2) occurs as two acrocentric chromosomes in another species, this pair may not undergo residual tetrasomy and therefore may have increased sequence divergence between homeologs. Therefore, species with few metacentric chromosomes may offer additional information regarding rediploidization. Even though Brook Charr has fewer metacentrics than other salmonids, all of the known residually tetrasomic homeolog pairs from the other salmonids have one homeolog present in a metacentric chromosome in Brook Charr (Table 2; n=8 metacentrics).

As metacentric formation is thought to be important for residual tetrasomy, the timing of fusion events may provide additional insight into the rediploidization process in salmonids. From the present study, it is interesting to note that many of the residually tetrasomic pairs have at least one homeolog involved in an ancient conserved fusion (Figure 4). The second homeolog varies more in its fusion partner across the lineage, or can be present as an acrocentric. For example, 9.2 of the 9.1/9.2 residually tetrasomic homeolog pair is fused with 17.1 in all assessed species (fused at F in Figure 4). In contrast, 9.1 varies more in its binding partner and sometimes is acrocentric in extant salmonids. Similarly, the residually tetrasomic 23.1 is fused with 24.1 in all assessed species except Lake Whitefish (10.2–24.1), whereas 23.2 is more variable and occasionally acrocentric. These ancient fusions may be informative about mechanisms that have prevented rediploidization in salmonids.

The fusion history of the other residually tetrasomic pairs are not as simple as the above two examples. Within a residually tetrasomic homeolog pair, it is not always the same homeolog in a metacentric fusion across species. This agrees with previous indications that only one of the homeologs must be bound in a metacentric to prevent rediploidization. For example, in Atlantic Salmon, 2.2 is metacentric and 2.1 is acrocentric, whereas in Brook Charr and all *Oncorhynchus* spp., 2.1 is the conserved metacentric (fused at E in Figure 4). Another example of this differing metacentric binding occurs for 20.1/20.2, where in Atlantic Salmon and Brook Charr 20.1 is in a conserved metacentric fusion and 20.2 is acrocentric, whereas in all *Oncorhynchus* spp. 20.2–16.2 is the conserved fusion. Therefore, even though these metacentrics may be required to retain residual tetrasomy, the homeolog bound in the metacentric chromosome can differ among the species. Further characterization of this will be facilitated with the production of high-density linkage maps for more species from salmonid genera outside of *Oncorhynchus* that are represented by only one species (e.g. *Salvelinus, Coregonus, Salmo)* or none (e.g. *Thymallus)*.

Although rediploidization of the salmonids may have generally occurred prior to the salmonid radiation (Lien et al. 2016), the rediploidization process has apparently continued since the speciation of Atlantic Salmon, at least in one homeologous pair (i.e. 6.1/6.2). This pair has rediploidized in *Salmo salar* as demonstrated by sequence similarity between homeologous chromosomes (see 18qa-1qa in Figure 3b in Lien et al. 2016) as well as suggested by the lack of identifiable isoloci in this pair (Lien et al. 2011). Conversely, in *Oncorhynchus*, this homeologous pair is residually tetrasomic as demonstrated by isoloci in Coho, Chinook, Chum and Sockeye Salmon (see Table 1 for references). This difference in rediploidization between Atlantic Salmon and the Pacific salmonids is shown from the orthologous relationships of chromosome arms characterized in Table 2.

It is interesting to note that one of the homeologous chromosome arms, 6.1, is fused in the center of one of the triple chromosome arm fusions specifically in Atlantic Salmon (Figure 4). Although it is known that residual tetrasomy requires at least one of the two homeologous chromosomes to be in a metacentric fusion, the effect of being in the middle of a triple chromosome arm fusion on rediploidization is not known. It is possible that this position could hinder homeologous pairing at meiosis. Regardless of this mechanism, this result indicates that the path to rediploidization differs for this chromosome pair between Atlantic Salmon and the evaluated Pacific salmonids. Information regarding residual tetrasomy (e.g. through haploid crosses) in additional maps from members of *Coregonus, Salvelinus, Thymallus* or other salmonid genera will be valuable to understand this process further in a broader range of genera.

### Fusions and inversions

Chromosomal rearrangements include chromosome fusions or fissions, region amplifications or deletions, segment inversions or non-homologous chromosome segment translocations (Rieseberg 2001). The characterization of the fusion events across all published salmonid maps (Figure 4) provides a new resolution of the exact identities of chromosome arms in the preduplicated genome that have fused together at different moments during the salmonid diversification. This demonstrates the stepwise process of generating the extant salmonid karyotypes, with fusions occurring at each step along the diversification process. Notably, for most salmonid species, most fusions are not ancestrally conserved, but rather occur individually within each species (Figure 4). It remains unclear why some species retain their high number of acrocentric chromosomes (e.g. Brook Charr, three species-specific fusions), whereas others do not (e.g. Pink Salmon, 17 species-specific fusions). Furthermore, this variation in numbers of species-specific fusions can occur between closely related species (e.g. Chum and Sockeye Salmon).

Inversions can occur when a segment of a chromosome is cut out by two breakpoints and then reinserted in the opposite orientation (Kirkpatrick 2010). Effects of inversions on fitness are highly unpredictable and vary across taxa. In general, they tend to reduce recombination rates at the site of the inversion, potentially playing an important role in speciation and local adaptation (Rieseberg 2001; Kirkpatrick 2010; Noor et al. 2001a; 2001b). For example, two inversions reduce recombination and maintain genetic differentiation between migratory and stationary ecotypes of Atlantic Cod *(Gadus morhua)*, preserving the co-occurrence of adaptive alleles within the migratory form (Kirubakaran et al. 2016) Additionally, lower recombination rates were observed in heterokaryotypic regions of Yellowstone Cutthroat Trout *(O. clarkii)* and Rainbow Trout hybrids compared to collinear regions (Ostberg et al. 2013). Recombination suppression may allow for conservation of fitness-related gene complexes that are locally adapted, or involved in reproductive isolation (Ostberg et al. 2013). Robertsonian rearrangements (e.g. fusions and fissions) have less of an effect on recombination rates than do rearrangements affecting synteny (e.g. inversions) (Rieseberg 2001; Ostberg et al. 2013).

### *MapComp:* Potential and limitations

By using the information from both identical and proximal marker pairing, MapComp solves the issue of low marker homology between reduced representation sequencing (RADseq)-based linkage maps generated with different protocols or restriction enzymes, or from relatively more distantly related species. Synteny is still required in order to pair proximal markers through the intermediate reference genome. Previously, polymorphic microsatellite markers highly conserved among salmonids have enabled exploration of salmonid chromosomal evolution by integrating across species and genera (Naish et al. 2013). Although RADseq-based linkage maps routinely provide an order of magnitude more markers than microsatellite maps with less effort, identical markers are not always abundant between species. Low marker homology among species has also hindered cross-species comparisons when using microsatellite-based genetic maps, for example when Coho Salmon was compared with Sockeye Salmon and Pink Salmon (Naish et al. 2013). As such, with the generation of additional high-density maps for the salmonids, the use of MapComp will continue to be highly useful in characterizing these relationships.

At its core, MapComp is similar to the approach used by Sarropoulou et al. (2008), in which EST-based markers from two species were aligned to a reference genome of a third species, to identify orthologous linkage groups. However, this earlier approach did not retain marker positions from original maps for plotting within an 0xford grid, and only provided the total number of markers found to correspond for each linkage group pair. Other cross-species map comparison approaches exist, for example cMAP (Fang et al. 2003), although these often require shared markers between maps. Another similar approach was used by Amores et al. (2011) for Spotted Gar *Lepisosteus oculatus*, where paired-end sequencing was performed on a single-digest then random shear library. The authors therefore obtained a larger amount of sequence near their marker allowing them to identify genes near the marker. Then the order of the identified genes was used to compare synteny of orthologs in assembled genomes such as humans *Homo sapiens* or Zebrafish *Danio rerio*. In contrast, MapComp works without prior knowledge of specific gene orthology, providing map comparisons at a much higher marker density without being restricted to coding regions. Another recent approach compared a linkage map for the European tree frog *Hyla arborea* with the genome of the western clawed frog *Xenopus tropicalis* and identified many syntenic regions (Brelsford et al. 2015). A recent approach in salmonids used a RADseq high-density linkage map for Chinook Salmon with the Atlantic Salmon reference genome to anchor Atlantic Salmon scaffolds to the Chinook Salmon linkage map when enough markers were present and the order was as expected (McKinney et al. 2015). Orthology relationships between Chinook Salmon and Atlantic Salmon have been characterized previously (Brieuc et al. 2014), and so the aligned scaffolds could then be classified as homologous, homeologous, or unsupported to further improve the anchoring of scaffolds onto the linkage map, and to identify potential genes around loci of interest (McKinney et al. 2015). MapComp is not meant to be used for RADseq based phylogenetic analysis, which requires identical markers for comparisons; this is rather performed using the direct marker approach with reciprocal best hit BLAST (Cariou et al. 2013; Pante et al. 2014).

MapComp is thus an easy solution to compare genetic maps in a way that is more tolerant of different library preparation protocols and phylogenetic distances. As shown here, MapComp is effective at finding orthology between chromosomes (Table 2), permitting the characterization of chromosomal rearrangements since whole genome duplication (Figure 4) and identifying putative structural rearrangements (Figure 5). This method will allow for the exploration of corresponding regions between species, such as regions harboring QTLs (Sarropoulou et al. 2008). Advances in genomics have resulted in many taxonomic groups having at least one species with a reference genome at some stage of assembly, providing the intermediate genome needed for this approach, and opening up this approach for a number of other taxonomic groups. MapComp is freely available at: https://github.com/enormandeau/mapcomp/

## Conclusions

We provide the most complete analysis to date of the chromosomal rearrangements that lead to the current chromosome conformations in salmonids using the newly developed MapComp method. This analysis permitted the integration of all high-density salmonid maps across the lineage, identifying the timing of fusions of all chromosomes, including those still undergoing residual tetrasomy in the characterized species. Large inversions were also identified using this method, including a pericentric inversion that has occurred after the ancestral fusion of two chromosome arms and potentially rearranged the position of 11 genes across a centromere. These analyses will be further refined through the continued availability of other high-density salmonid maps, and can provide insights into the chromosomal evolution in both salmonids and other taxa.

## Acknowledgements

This work was funded by a Fonds de Recherche Nature et Technologies (FRQNT) research grant awarded to Céline Audet, Louis Bernatchez and Nadia Aubin-Horth and a grant from the Société de Recherche et de Développement en Aquaculture Continentale (SORDAC) awarded to Céline Audet and Louis Bernatchez. We would like to thank the staff of the Institut des Sciences de la Mer de Rimouski (ISMER), Pierre East from Pisciculture de la Jacques-Cartier, the Reseau Aquaculture Quebec (RAQ), and Pierre Dubé from SORDAC. We also express our thanks to G. Côté, G. Legare, C. Rougeux and B. Boyle from the Institut de Biologie Integrative et des Systemes (IBIS) for help with DNA extraction and library preparation. We also thank A-M Dion-Côté, E Rondeau, M. Limborg, two anonymous Reviewers and D. Huchon for valuable comments on the manuscript. BJGS was supported by an NSERC postdoctoral fellowship. Thanks to the members of the Bernatchez Laboratory for discussion and support.

## Supporting Information

**Additional File S1. Mapping software parameters and bioinformatics pipeline overview.** STACKS and JOINMAP parameters and an outline of bioinformatics steps used to generate the Brook Charr linkage map.

**Additional File S2. Descriptive statistics for Brook Charr female map**.

**Additional File S3. Brook Charr female linkage map**. Includes species name, linkage group, cM position, marker name and sequence.

**Additional File S4. Putative genes within the 9.2-17.1 pericentric inversion**. Genes annotated in the Atlantic Salmon genome within the putative inverted region in the Pink, Chum and Sockeye Salmon clade.

**Additional File S5. Brook Charr and all other species in Oxford grids.** Oxford grids comparing Brook Charr to other species using the Rainbow Trout genome as the reference intermediate, including: 1) Lake Whitefish, 2) Atlantic Salmon, 3) Rainbow Trout, 4) Coho Salmon, 5) Chinook Salmon, 6) Pink Salmon, 7) Chum Salmon, 8) Sockeye Salmon.

**Additional File S6. Distribution of paired marker distances**. Frequencies of distances between marker pairs in various comparisons via MapComp. As can be seen, more closely-related species have more homologous markers (0 distance between marker pairs), whereas the more distantly related comparisons have fewer homologous markers.

## Code and Pipeline Availability

MapComp: https://github.com/enormandeau/mapcomp/

Collecting and formatting available salmonid maps: https://github.com/bensutherland/2016_ms_sfonmap

RADseq workflow: http://gbs-cloud-tutorial.readthedocs.org

STACKS workflow: https://github.com/enormandeau/stacks_workflow

